# The Mechanism Insight into Bacterial Degradation of Pentachlorobiphenyl

**DOI:** 10.1101/2024.01.18.576235

**Authors:** Lei Ji, Xiaoyu Chang, Leilei Wang, Xiaowen Fu, Wenkai Lai, Liwen Zheng, Qi Li, Yingna Xing, Zhongfeng Yang, Yuyao Guan, Fenglong Yang

## Abstract

Bacterial degradation mechanism for high chlorinated pentachlorobiphenyl (PentaCB) with worse biodegradability has not been fully elucidated, which could limit the full remediation of PCBs-combined pollution. In this research, using enzymatic screening method, a new PentaCB-degrading bacterium *M. paraoxydans* that has not been reported was obtained. The characteristic of its intracellular enzymes, proteome and metabolome variation during PentaCB degradation were investigated systematically. The results showed that PentaCB (PCB101, 1 mg/L) degradation rate could arrive 23.9% within 4 h till complete degradation within 12 h. The intracellular enzyme compound was optimally active at pH 6.0. The 12 up-regulated characterized proteins involved ABC transporter substrate-binding protein, translocase protein TatA and signal peptidase I (SPase I) indicated that functional proteins for PentaCB degradation were present both in the cytoplasm and outer surface of cytoplasmic membrane. There were also 5 differential metabolites strongly associated with above proteins in which the up-regulated 1, 2, 4-benzenetriol was enriched into the degradation pathways of benzoate, chlorocyclohexane, chlorobenzene and aminobenzoate. Bacterial degradation of PentaCB necessitates transmembrane transport, energy consumption, protein export, biofilm formation and quorum sensing. These findings hold significant theory and application value for PCBs biodegradation.

## 1. INTRODUCTION

Industrialization and intensive use of chemical substance polychlorinated biphenyls (PCBs) are contributing to persistent and toxic environmental pollution (Megharaj et al., 2011; Furukawa and Fujihara, 2008). PCBs in submarine canyon sediments are several times higher than those in sediments from continental shelfs (Sevastyanov et al., 2020; Zhou et al., 2022). Removal of PCBs from contaminated environments has been primarily performed by incineration at high temperatures, which may entail additional energy consumption and greenhouse effect (Stella et al., 2017). Mild industrial conditions are beneficial for preventing modification of original heat-sensitive substrates and generation of adverse by-products (Ji et al., 2014). Biological PCBs clean-up methods have a considerable potential as a well-established and cost-effective strategy among methods employed for waste treatment and environmental remediation at ambient temperature (Šrédlová and Cajthaml, 2022). Microbial degradation of PCBs is highly strain-dependent (Furukawa and Matsumura, 1976; Han et al., 2023) and screening for effective microbes is a time-consuming process, particularly when performed on a large scale. While several studies have examined bacteria that oxidatively degrade PCBs, encompassing both gram-negative and gram-positive genera (Megharaj et al., 2011; Furukawa and Fujihara, 2008), there still remains a pressing need for highly efficient screening methods and bacteria that can effectively degrade PCBs for practical application.

The majority of bacterial aromatic degradation is due to a series of enzymatic reactions, including the formation of diols, incorporation of oxygen atoms into the aromatic rings, and ring cleavage (Lee et al., 2016; Suenaga et al., 2009). Enzymes catalyze metabolic transformations under mild conditions with high catalytic rate and reaction selectivity, which could be realized in a shortened time (Ji et al., 2019). The screening speed of functional microbes could be much higher through bio-catalysis using enzymatic method. Moreover, bioconversion catalyzed by enzymes from microbes improves the bioavailability and the consequent biodegradation without prior desorption, which could more truly reflect the microbial degradation potential.

Commercial PCB mixtures are mostly tri-to hexachlorinated ones of 209 congeners (Furukawa and Fujihara, 2008). The di– and trichlorobiphenyls are the most readily degraded congeners (Gilbert and Crowley, 1998). Biodegradability decreases with increased number of chlorines, and PCB congeners with chlorine at position 2, 2′-(double ortho-substituted congeners) are poorly degraded (Furukawa and Fujihara, 2008). Wang et al. (2013) showed that the concentration of the most toxic dioxin-like PCBs (DL-PCBs) in surface soils ranged from 1.4 μg·kg^-1^ to 7.4 μg·kg^-1^ in Gudao town with chemical plants around, Dongying City, exceeding the Canadian soil environment quality guidelines. Tetrachlorobiphenyl (TetraCB) and pentachlorobiphenyl (PentaCB) were the major homologues, together accounting for more than 80% of the total DL-PCBs (Wang et al., 2013). Sanli et al. (2023) found that in industrial, urban and semi-rural areas of Bursa, Turkey, total PCBs content in spring surface soils reached 8.4∼9.6 μg·kg^-1^, 2.9∼4.1 μg·kg^-1^, 2.5∼4.9 μg·kg^-1^, respectively, and was similarly dominated by TetraCBs and PentaCBs (Sanli et al., 2023). Previous studies about biodegradation of PentaCB are most involved in cometabolism with biphenyl or PCBs mixture (e.g. Aroclor mixtures) (Čvančarová, et al., 2012). Bacterial degradation mechanism of PentaCB on protein and metabolite levels has not been fully elucidated. Incomplete biological information regarding the cellular responses restricts progress in the bioremediation process (Zhao and Poh, 2008).

In this study, an enzymatic screening approach was employed to acquire highly efficient bacteria capable of degrading PentaCB. Subsequently, the enzyme characteristics of the most potent bacterium, *Microbacterium paraoxydans*, were investigated to elucidate the optimal conditions, given the scarcity of reports on the *Microbacterium genus*’ PCBs degradation capacity. Additionally, the cellular responses of *M. paraoxydans* to PentaCB stimuli were examined at the proteome and metabolome levels using quantitative analysis of the proteome through TMT and untargeted metabolomics via gas chromatography-mass spectrometry (GC-MS). Consequently, a novel observation was made regarding the comprehensive alterations in differential proteins and metabolites of the *Microbacterium genus* throughout the biodegradation process of PentaCB.

## 2. MATERIALS AND METHODS

### 2.1 Preparation of Bacterial Intracellular Enzymes for PentaCB Degradation

LB medium (Table 1S) was used for strain activation at 30°C and 150 rpm. Induction medium (Table 2S, pH7.0) containing 1.0 mg·L^-1^ PCB101 (2, 2’, 4, 5, 5’-PCB) was used for PentaCB degrading enzymes production at 30°C and 150 rpm. Seed liquid was introduced into the induction medium and incubated. When the value of OD_600_ showed that the culture reached the end of the exponential phase, it was harvested by centrifugation at 4 °C and 8,000 rpm for 5 min. The cells were subsequently re-suspended in a 10 mM potassium phosphate buffer (4 mM DDT, pH7.0) and sonicated for 17 min at 320 W using a sonicator (SM-650D, Nanjing Shunma), with a sonication time of 3 s and quench time of 5 s (Ji et al., 2019). Following ultrasonic treatment, the suspension was centrifuged and the resulting supernatant was obtained as the PentaCB degrading enzymes.

### 2.2 Efficiency Determination of PentaCB Degradation Enzymes

Each reaction system was inoculated with different load of above PentaCB degrading enzymes and reacted under pH4.0-8.0 separately. The control samples contained no PentaCB degrading enzymes. Harvests were carried out at different time points and the rate of PentaCB degradation was determined by GC-MS as followed and then calculated.

PentaCB was extracted using Dionex 200 Accelerated Solvent Extraction (ASE) system and then subjected to GC-MS analysis utilizing the 450-GC, 240-MS ion trap detector from Varian (Walnut Creek, CA), as previously reported (Grigorakis and Drouillard, 2018). All experiments were performed in triplicate, and the presented results represent the mean values and standard deviations.

### 2.3 Sample Preparation and Quantitative Analyses in Proteomics and Metabolomics

Cells of *M. paraoxydans* (CGMCC No.15836) were cultured until the late stationary phase in the induction medium comprising PCB101, and subsequently harvested by centrifugation at 4 °C and 8,000 rpm for 5 min. The precipitation cells were then subjected to three rounds of washing with an equal volume of ice-cold 0.2 M sucrose, followed by a final wash with ice-cold methanol. Meanwhile, cells without induction by PCB101 were prepared as controls (CK) under identical conditions. Proteomics and metabolomics quantification were performed at Novogene Bioinformatics Technology Co., Ltd. (Beijing, China). Detailed sequencing procedures are presented in supplementary materials (S1, S2).

## 3. RESULTS

### 3.1 Enzymatic Screening for Bacteria Capable of Degrading PentaCB

Intracellular enzymes from six biphenyl metabolic bacteria (Figure 2A) were collected separately, and then subjected to incubation with 1 mg·L^-1^ PCB101 at 30°C and 150 rpm for 12 h. All of the strains exhibited the capability to degrade PCB101 (Figure 2A). Notably, the intracellular enzymes from *M. paraoxydans* demonstrated the highest efficiency, decomposing 100% of PCB101. *Arthrobacter phenanthrenivorans* and *Bacillus* sp. ECO-12 followed with degradation rates of 39.6% and 30.8%, respectively.

**Figure 1.**
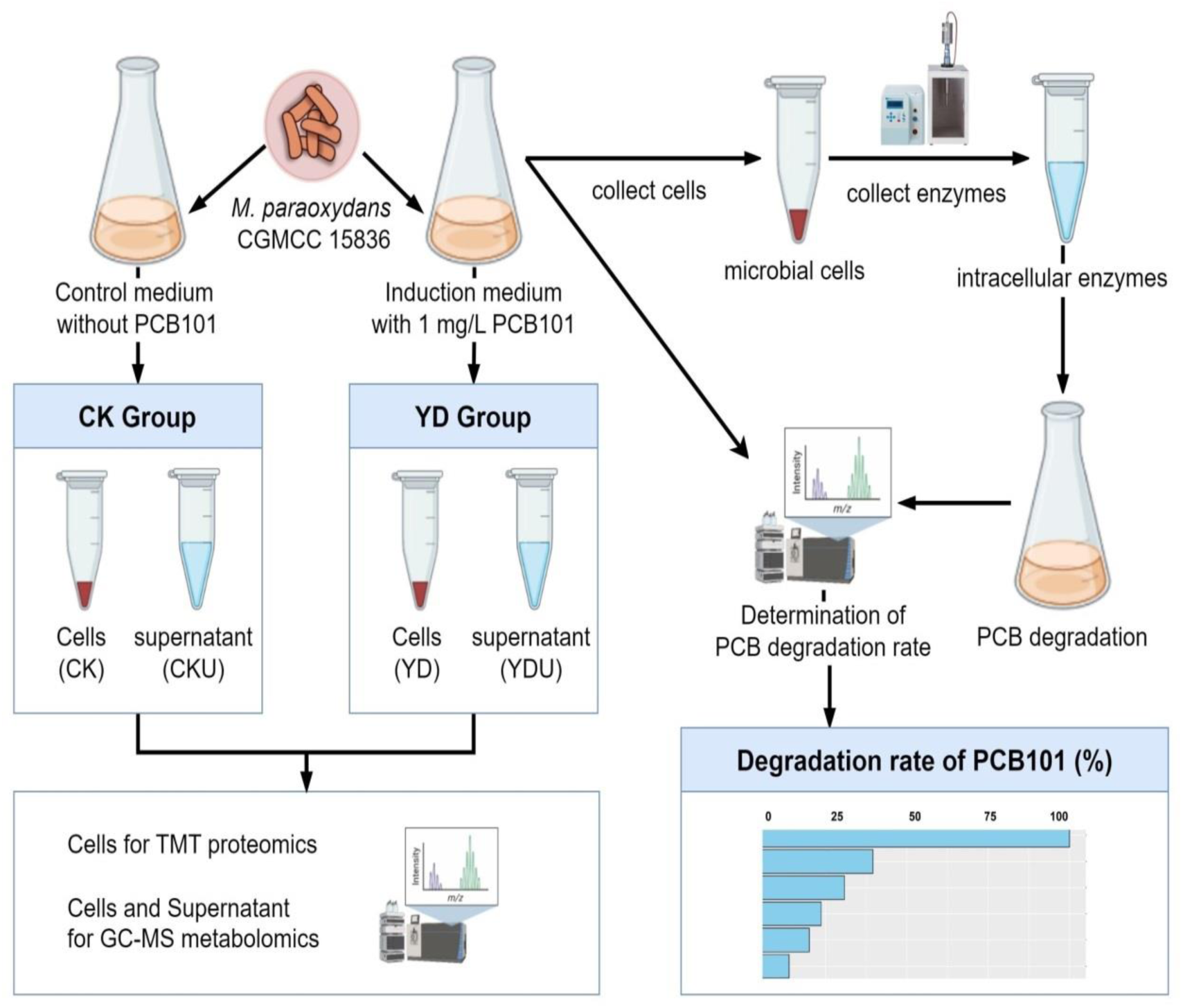
Overview of the workflow. (CK: Cells without induction; YD: Cells induced by PentaCB; CKU: The supernatant of cells without induction; YDU: The supernatant of cells induced by PentaCB.)

**Figure 2.**
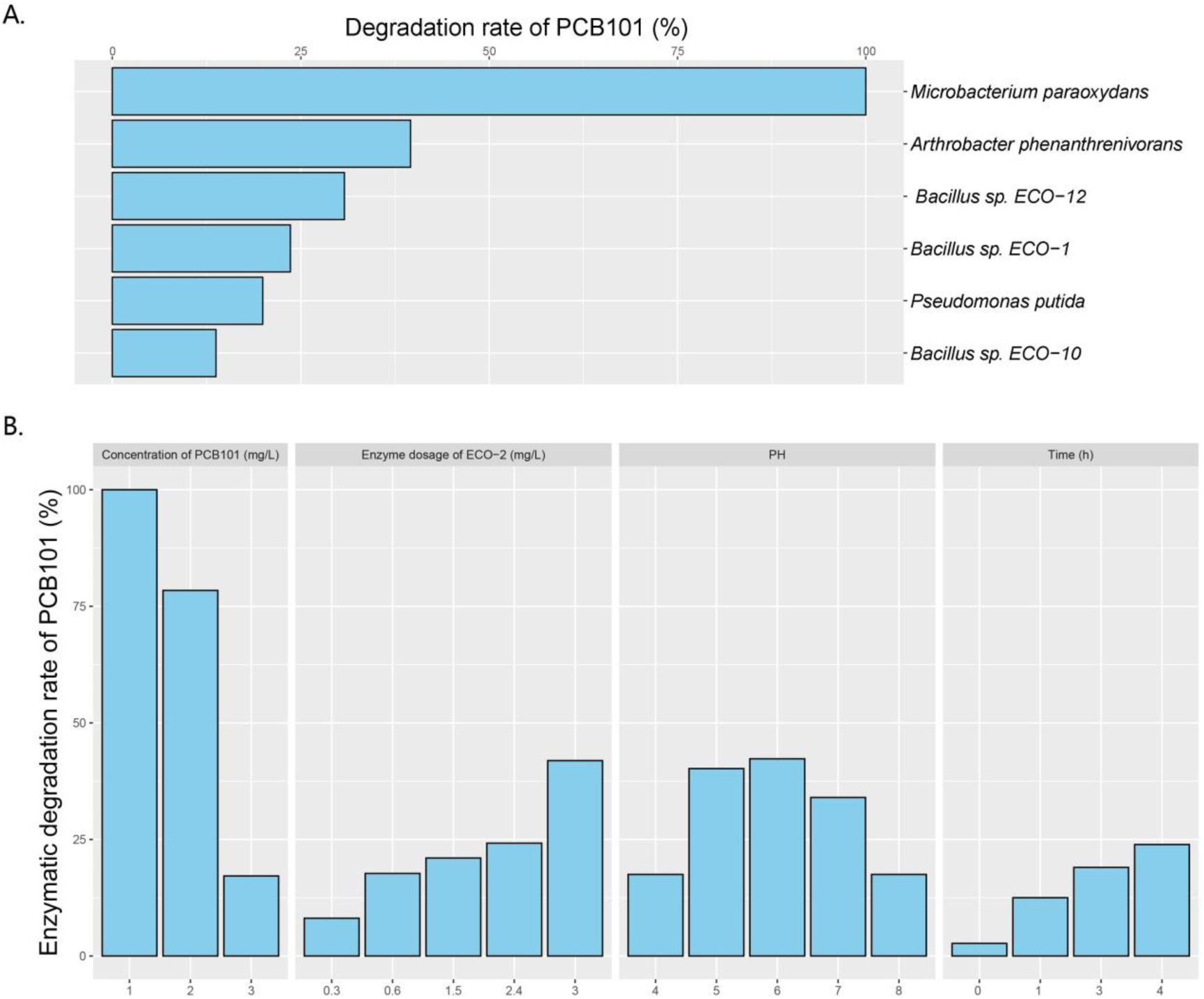
PentaCB-degrading capabilities by intracellular enzymes from biphenyl metabolic bacteria. (A) Enzymatic reaction with 1 mg·L^-1^ PCB101 for 12 h. (B) Enzymatic reaction under different concentrations of PCB101, enzyme dosages, pH levels and time courses.

Wang et al. (2017) applied an enzymatic method instead of the shaking culture one to analyze the microbial metabolism capability of petroleum hydrocarbons, and the testing time was substantially reduced. In our study, it was further verified that this means was more efficient in distinguishing functional bacteria capable of pollutant degradation, which was valuable for surveying environmental resources.

### 3.2 Enzymes Characterization for PentaCB Degradation

The characteristics of intracellular enzyme from bacterium *M. paraoxydans* were measured for the optimal operating conditions during PentaCB degradation. It was showed that PCB101 degradation rate decreased to 78.4% and 17.2% respectively following its progressively increased concentration (Figure 2B). The addition of enzyme enhanced the degradation efficiency. Similarly, reducing the enzyme dosage by half, the degradation rate of PCB101 maintained directly proportional to the enzyme content. Within 2 h and 4 h, the enzymatic degradation efficiency of PCB101 reached 12.5% and 23.9%, respectively. This indicates the potential to significantly shorten the high-throughput screening process for functional microbes. The optimal pH for enzyme activity was found to be 6.0, with stability observed over a broad pH range of 4.0 to 8.0.

### 3.3 Proteomic Insights during PentaCB Degradation

Quantification of proteome changes under PentaCB stress were analyzed, resulting in a total of 885,055 matched spectra to known ones. Additionally, 17,460 unique peptides were generated, and a total of 2,231 proteins were detected.

To obtain more detailed functional information about the quantified and identified proteins, a Venn diagram was used to compare the Gene Ontology (GO), clusters of orthologous groups (COG), Kyoto encyclopedia of genes and genomes (KEGG), and InterPro (IPR) domain annotations (Figure 3A). Based on GO terms, a total of 1,354 proteins were annotated across three categories: biological process (BP), cellular component (CC), and moleculr function (MF). The most commonly annotated category under BP term was the oxidation-reduction process including 161 proteins. Under the CC term, 123 proteins were annotated as being associated with membranes. In the MF category, 185 proteins were annotated as ATP binding. Using the COG functional classification, a total of 2,026 proteins were annotated. The main functional classification was amino acid transport and metabolism, which involved 261 proteins. Furthermore, 1,955 proteins were annotated in different KEGG pathways. The metabolism pathway with 1,178 proteins identified had the highest number of annotated proteins. Simultaneously, 1,974 proteins were annotated with different IPR terms. The most common IPR-annotated proteins included the AAA+ ATPase domain (90 proteins), the ABC transporter-like domain (72 proteins), and the ABC transporter type 1 transmembrane domain MetI-like (40 proteins).

**Figure 3.**
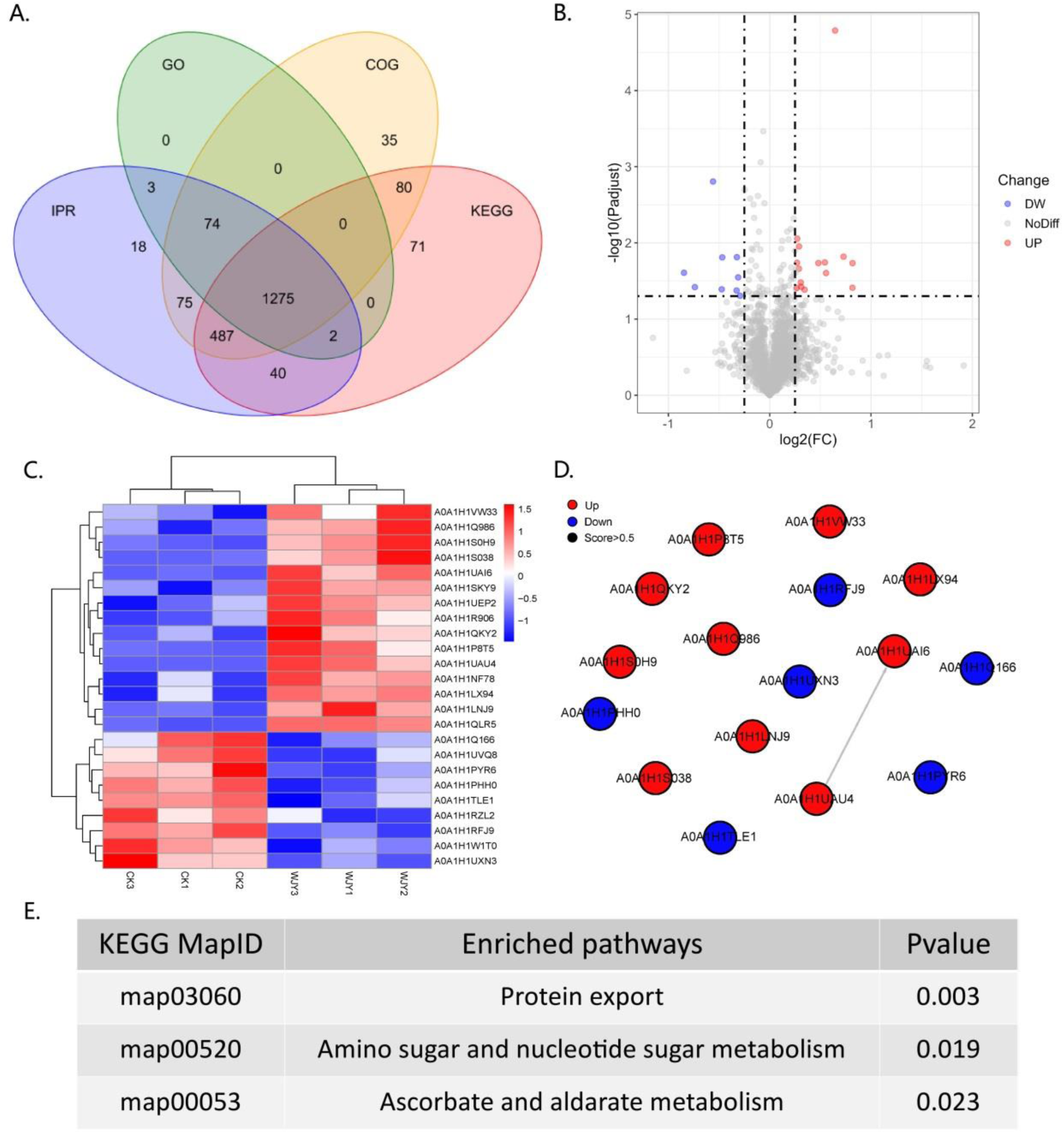
Insights from Proteomic Analysis of PentaCB Degradation by *M. paraoxydans*. (A) Venn diagram of function annotation based on enriched GO terms, GOG functional classification, KEGG pathways and IPR terms. (B) Volcano plot of differential proteins between YD and CK groups. Each dot represents one protein. Red dots represent the significantly up-regulated proteins and green dots represent the significantly down-regulated proteins. Gray dots represent no significantly differential proteins. (C) Clustering heat map of differential proteins. (D) PPI network. (E) KEGG enriched pathways of differential proteins (p<0.05).

Here, there were 24 differential proteins detected in which 15 up-regulated and 9 down-regulated proteins were included (Figure 3B). Among the up-regulated proteins, 12 characterized proteins were identified, including carbohydrate ABC transporter substrate-binding protein, L-alanine-DL-glutamate epimerase, peptidase inhibitor I9, galactokinase, DNA-binding transcriptional regulators (Lrp family, PadR family, Lrp/AsnC family), putative adhesion proteins, Sec-independent protein translocase protein TatA, bifunctional protein GlmU, sugar phosphate permease, and signal peptidase I (SPase I) (Figure 3C). Based on MF category, these up-regulated proteins were involved in serine-type peptidase activity, galactokinase activity, sequence-specific DNA binding, and protein transporter activity. Besides, the differentially up-regulated proteins were enriched in pathways of protein export, amino sugar and nucleotide sugar metabolism, bacterial secretion system, galactose metabolism, and quorum sensing (Figure 3E).

### 3.4 Metabolomic Analysis during PentaCB Degradation

Metabolic profiling was performed to gain insights into the efficient degradation mechanism of PentaCB by *M. paraoxydans*. A total of 283 metabolites were identified and screened for differential accumulation. The top five differential metabolites distinguishing YD from CK (Figure 4) were determined based on VIP (variable importance in projection) score > 1.0, absolute value of log_2_ (fold change) > 1.2, and p < 0.05. Four metabolites including 1, 2, 4-benzenetriol (C_6_H_6_O_3_), octadecanol, malonamide, citraconic acid (C_5_H_6_O_4_) showed significant up-regulation while hexadecane exhibited significant down-regulation (Figure 4B, C). Differential metabolites were enriched in pathways of benzoate degradation, chlorocyclohexane and chlorobenzene degradation, and aminobenzoate degradation (Figure 4D). The metabolic pathways were significantly enhanced in cells induced by PentaCB. The enriched pathways of benzoate degradation and aminobenzoate degradation were observed only in induced cells samples.

**Figure 4.**
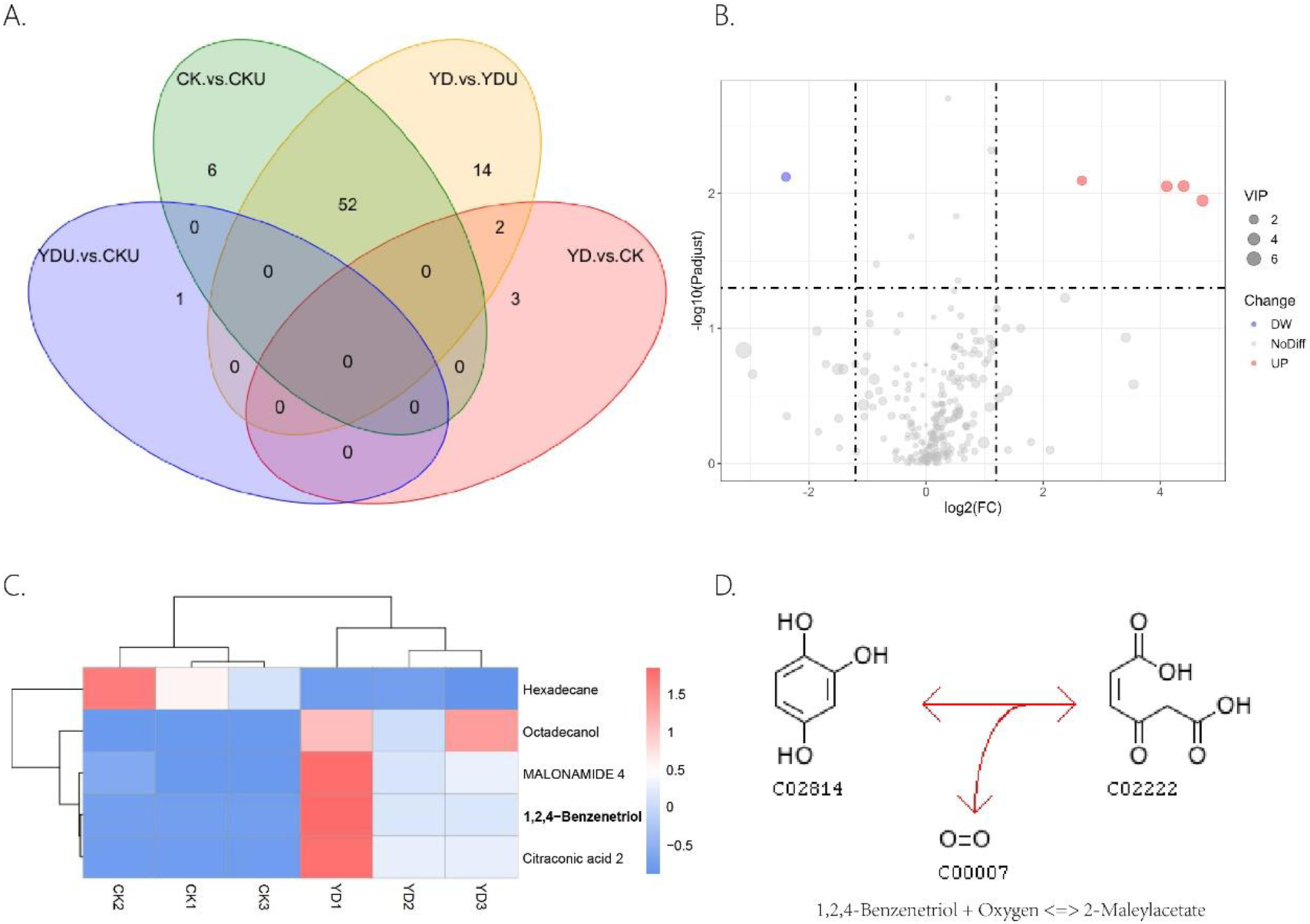
Insights from Metabolomic Analysis of PentaCB Degradation by *M. paraoxydans*. (A) Venn diagram for multigroup comparison of differential metabolites. (B) Volcano plot of differential metabolites between YD and CK groups. Each dot represents one metabolite. Red dots represent the significantly up-regulated metabolites and green dots represent the significantly down-regulated metabolites. Gray dots represent no significantly differential metabolites. The dot size represents the value of the VIP score. (C) Clustering heat map of differential metabolites. (D) KEGG enriched pathway of differential metabolite 1, 2, 4-benzenetriol. It reacts with oxygen to produce 2-maleylacetate. This reaction (R03891) is a constituent of the chlorocyclohexane and chlorobenzene degradation pathway (rn00361). CK: Cells without induction; CKU: The supernatant of cells without induction; YD: Cells induced by PentaCB; YDU: The supernatant of cells induced by PentaCB.

### 3.5 Multi-omics Analysis of PentaCB Degradation

Through a combined analysis of differential proteome and metabolome, it was found that the 5 most influential metabolites were strongly associated with the 24 differentially abundant proteins that distinguished YD from CK during degradation process of PentaCB (Figure 5).

**Figure 5.**
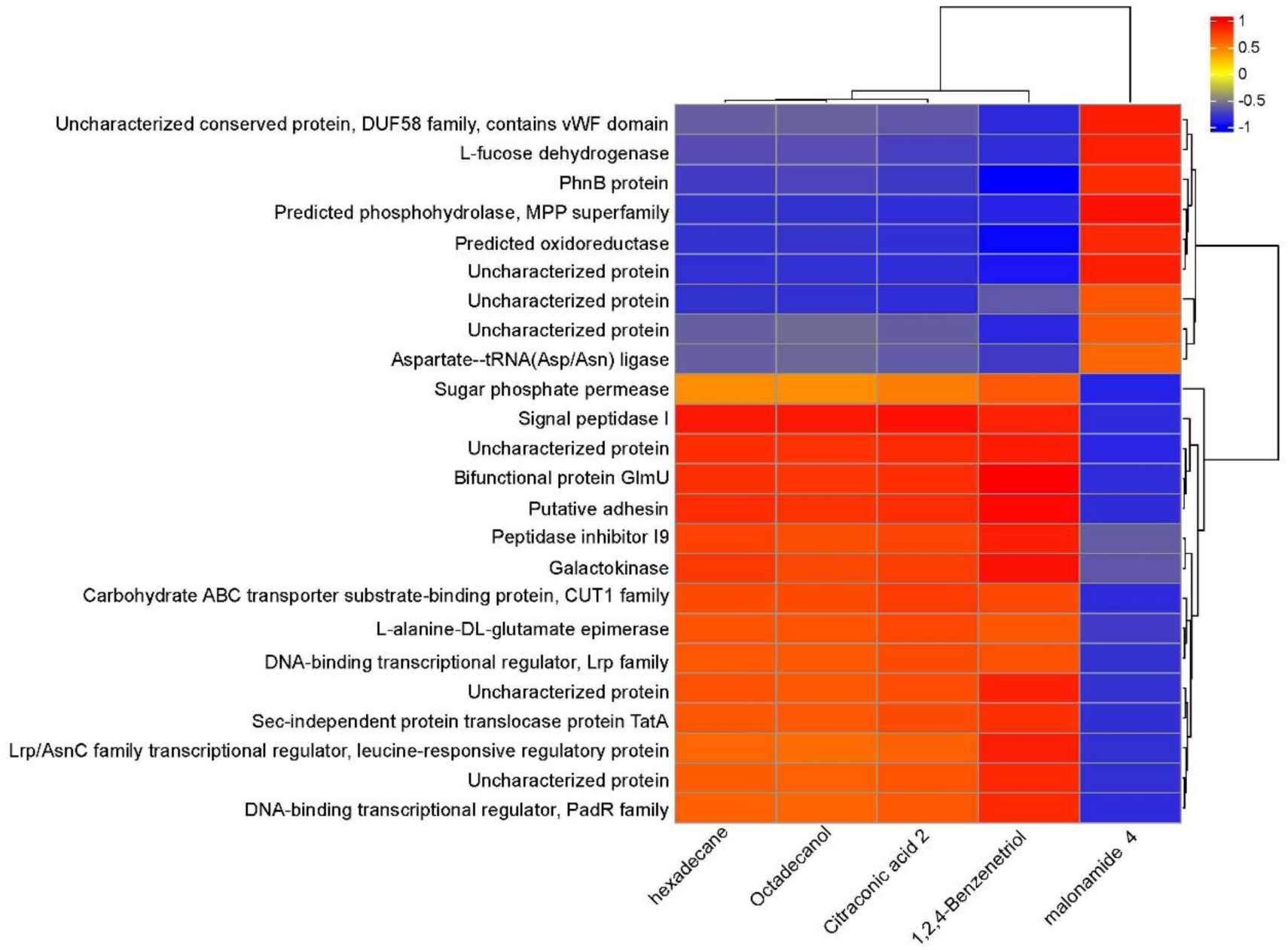
Correlation between differential proteins and metabolites during PentaCB Degradation by *M. paraoxydans*. The redder the color, the stronger the positive correlation, and the bluer the color, the stronger the negative correlation.

**Figure 6.**
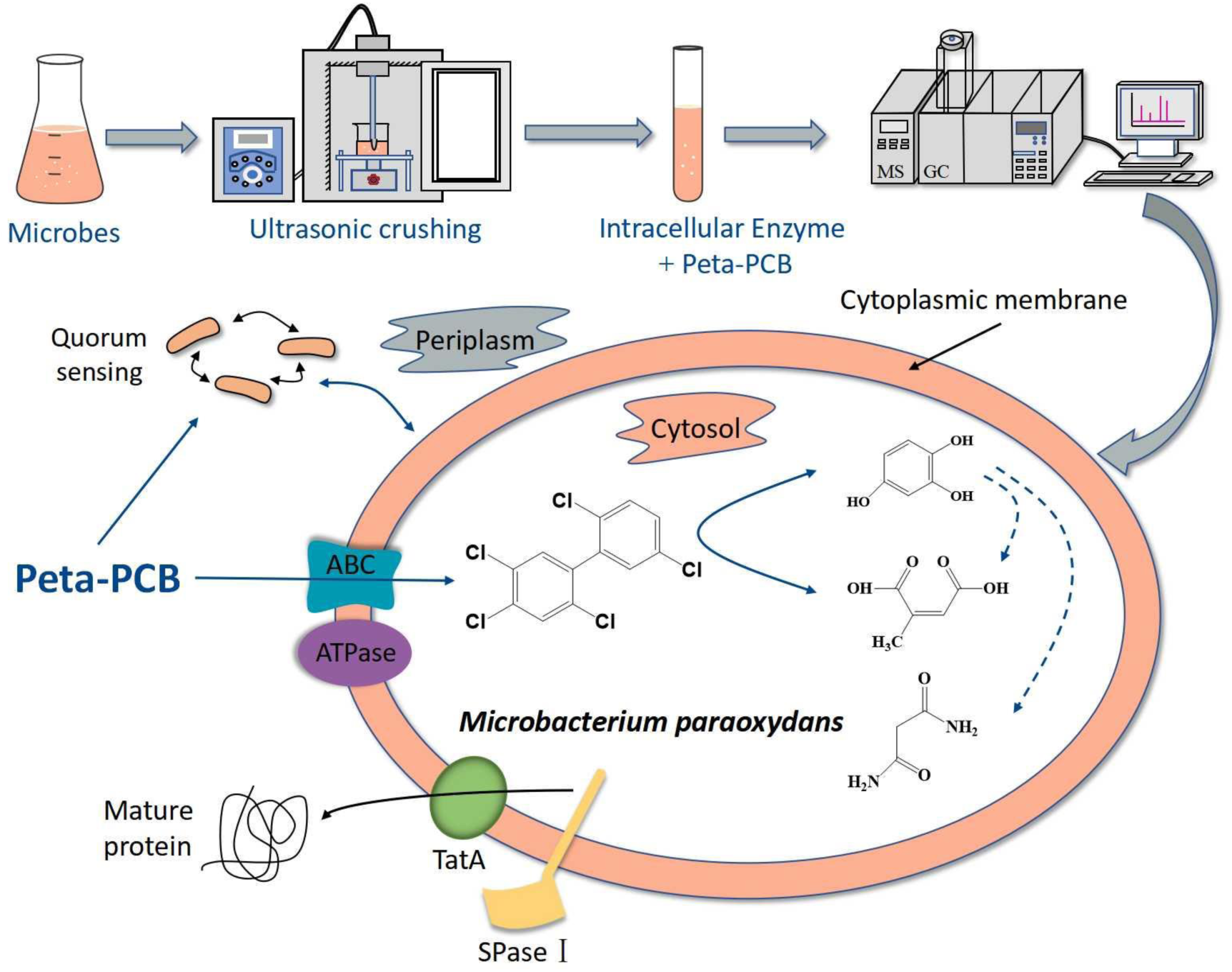
Hypothesized Mechanism of PentaCB Degradation by *M. paraoxydans*.

## 4. DISCUSSION

### 4.1 PCBs-Degrading Strains and Capabilities

Several genera with reported PCBs-degrading ability include *Rhodococcus, Bacillus, Pseudomonas, Achromobacter,* and *Burkholderia* (Field and Sierra-Alvarez, 2008; Vasilyeva and Strijakova, 2007; Stella et al., 2017; Han et al., 2023). Gilbert and Crowley (1997) observed that resting cells of carvone-induced *Arthrobacter* sp. B1B degraded 26 major components of Aroclor 1242, including 14±6% of PCB101 within 15 h, and noted that the more highly chlorinated congeners (such as tetra– and pentachlorinated ones) were transformed at a slower rate compared to the di– and trichlorobiphenyls. There were also 14 bacteria strains co-metabolizing biphenyl and PCBs isolated from environmental samples, belonging to the genera *Pseudomonas* and *Rhodococcus* (Takahito et al., 2016). And *Pseudomonas extremaustralis* ADA-5 can utilize decachlorobiphenyl (DCB) as the unique carbon source and degrade 9.75% DCB within 336 h (López et al., 2021). In previous studies, the white-rot fungi *Pleurotus ostreatus* was the most efficient known organism in PCBs degradation, decomposing 99.6% of low-chlorinated biphenyls (commercial mixture Delor 103 with 2.6% PentaCB) after 42 days and having the capability of breaking down penta– and hexachlorinated biphenyls (Čvančarová et al., 2012). For contaminated soil, *P. ostreatus* achieved a 50.5% removal of PCBs from the rhizosphere of dumpsite soils after 12 weeks of treatment, however, significant PCBs concentration reduction was not observed within the first 6 weeks of incubation in highly contaminated soil. Both *P. ostreatus* and *Irpex lacteus* were capable of efficiently degrading a wide range of aromatic organopollutants through the secretion of oxidative enzymes with low substrate specificity in the process of PCBs biotransformation (Stella et al., 2017). Resuscitating viable but non-culturable (VBNC) bacteria could provide huge candidates for obtaining high-efficient PCBs degraders. Lin et al. (2022) utilized resuscitation-promoting factor (Rpf) to screen a resuscitated strain *Streptococcus* sp. SPC0 from PCBs (tri– and tetrachlorinated ones) contaminated soil and found that it degraded 88.6% of total PCBs within 84 h. Han et al. (2023) used the similar methods to screen a strain *Bacillus* sp. LS1 under aerobic condition that degraded 59.6% and 50.1% of total PCBs (tri– and tetrachlorinated ones) with the concentrations of 10 mg/L and 20 mg/L within 84h, respectively. In this research, we report the species *M. paraoxydans* exhibiting efficient ability of PentaCB degradation for the first time to our knowledge. *M. paraoxydans* and its intracellular enzymes showed the application value in the remediation of highly toxic and poorly degradable PCBs pollutants.

**Table 1.**
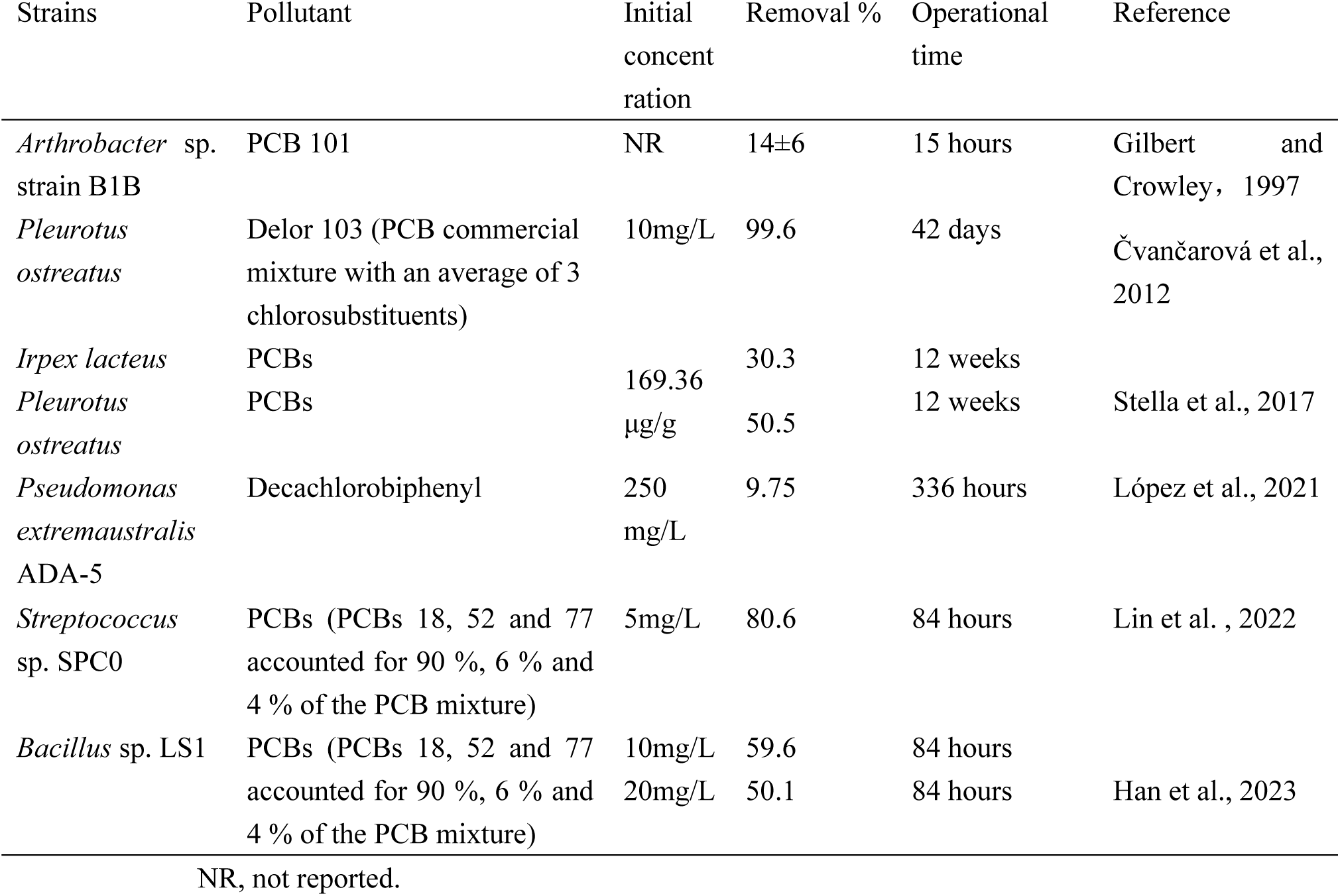
Overview of PCBs-degrading strains and capabilities.

### 4.2 Proteomic Dynamics in PentaCB Degradation

Functional enzymes of prokaryotic bacteria involved in organopollutants degradation were mainly intracellular (Kamei et al., 2006). The intake process of pollutants was considered as a rate-limiting step. For example, the uptake of long-chain alkanes is the first step in its utilization by many gram-negative bacteria (Liu et al., 2023). Using iTRAQ and PRM techniques, Jiang et al. (2021) demonstrated that the synthesis of peptide ABC transporter substrate-binding protein increased during the adsorption, uptake, and degradation of fluoranthene by *Bacillus cereus* in the presence of Tween 20. In our study, it was also found that the expression of carbohydrate ABC transporter substrate-binding protein involved in transmembrane transport was up-regulated during PentaCB degradation.

Besides, as it was known, ABC transporters primarily utilized ATP binding and hydrolysis to transport different substrates into cells (Jiang et al., 2021). The transmembrane transport of fluoranthene in *B. cereus* required transmembrane proteins and energy (Jiang et al., 2021). In our study, GO and IPR functional analysis also showed transmembrane transport involved energy consumption in *M. paraoxydans* for PentaCB degradation, which could be categorized into active transport.

Majority proteins in the cytosol of gram-positive bacteria were exported through the two systems, the general secretion (Sec) system or the twin-arginine translocation (Tat) system (Sargent et al., 1998; Berks et al., 2000; Palmer and Berks, 2012). The latter was known for its ability to export completely folded proteins across the cytoplasmic membrane (Freudl, 2013). In this study, as one of the integral membrane proteins of Tat translocase, the expression of Sec-independent translocase protein TatA was differentially up-regulated (Figure S1) and enriched in the pathway of protein export and bacterial secretion. Arauzo-Aguilera et al. (2023) reported recently protein YebF could be exported to the periplasm at high level through Tat pathway.

Moreover, another differentially up-regulated protein, the unique serine endoprotease SPase I, was essential for bacterial viability (Szałaj et al., 2023) and required by bacterial protein translocation pathways (Sec or Tat) for releasing translocated preproteins from the membrane during their transport from the cytoplasmic site of synthesis to extracytoplasmic locations (Auclair et al., 2012). In the case of *M. paraoxydans*, carbohydrate ABC transporter substrate-binding protein, TatA and SPase I (Figure S1) were together up-regulated, indicating that bacterial enzymes involved in PentaCB degradation were distributed both in the cytoplasm and at the outer face of the cytoplasmic membrane.

Additionally, the differentially up-regulated proteins from *M. paraoxydans* under PCB101 stress were enriched in one pathway of quorum sensing, which suggested it played a role in promoting PentaCB degradation. As reported, gram-positive bacterium typically employed quorum sensing communication circuits mediated by processed oligopeptides to regulate various physiological activities, including symbiosis, movement, and biofilm formation (Jiang et al., 2021; Alonso-Echanove et al., 2001). At the same time, another up-regulated protein in our study, the bifunctional protein GlmU, was reported to be related to biofilm production in *E. coli*, *Staphylococcus epidermidis,* and *Staphylococcus aureus* (Burton et al., 2006; Suman et al., 2011; Sharma et al., 2016). GlmU was also found critical for biofilm formation in *Mycobacterium smegmatis* under alkylating stress (Somma et al., 2019).

### 4.3 Cellular Responses and Pathways in PentaCB Degradation

Bacteria responses to environmental changes were also analyzed by metabolomics here (Eguchi et al., 2017; Zhou et al., 2017). Previous studies on PCBs bioremediation by white-rot fungi *P. ostreatus* and *I. lacteus* detected transformation products such as hydroxylated and methoxylated PCBs, chlorobenzoates, and chlorobenzyl alcohols (Stella et al., 2017). However, in this study, chlorinated derivatives of hydroxy– and methoxy-biphenyls were not detected among the differential metabolites. Instead, pathways related to chlorocyclohexane and chlorobenzene degradation, benzoate degradation, and aminobenzoate degradation were enriched.

The accumulation of intermediate PCBs metabolites was found to be minimal during degradation processes (Čvančarová et al., 2012). While previous research detected methoxylated metabolites but no hydroxylated compounds (Kamei et al., 2006b), this study identified a significantly up-regulated metabolite, 1, 2, 4-benzenetriol, which is a benzene metabolite (Sommers and Schiestl, 2006). Additionally, citraconic acid, another up-regulated metabolite, is associated with pyrimidine, purine, glutathione, and cysteine and methionine metabolism (Eguchi et al., 2017). Glutathione levels have been shown to decrease during PCBs treatment (Lai et al., 2010; Nomiyama et al., 2023; Zarerad et al., 2023), and PCBs metabolites can react with glutathione to form hydroquinone adducts of PCBs (Amaro et al., 1996). The increased octadecanol may enhance the hydrolytic action of phospholipases, which played a critical role in cell proliferation and metastasis (Bedia et al., 2015; Foster and Xu, 2003). On the other hand, hexadecane, which was significantly down-regulated, is used as a reaction medium for catalytic hydrodechlorination (Murena and Schioppa, 2000). These findings suggest that *M. paraoxydans* has the inherent ability to degrade PentaCB. The differential metabolites observed in *M. paraoxydans* cell samples, rather than the supernatant, indicate the cellular response to PentaCB stress, involving aromatic moiety decomposition, hydrodechlorination, and cell proliferation and metastasis.

## 5. CONCLUSION

In this research, an efficient enzymatic screening method helped to obtain a new PentaCB-degrading bacterium *M. paraoxydans*. Further, proteomic and metabolomic sequencing and deep analyses were employed to investigate the aerobic bacterial degradation mechanism of PentaCB. The differential proteins and metabolites together revealed that PentaCB transport into cytoplasm required transmembrane proteins such as ABC transporter substrate-binding protein and energy expenditure. And functional bacterial proteins for PentaCB degradation were distributed both in the cytoplasm and outer surface of cytoplasmic membrane. Protein export, bacterial secretion, biofilm formation and quorum sensing also played crucial roles simultaneously during bacterial degradation of PentaCB (Figure S1). These findings will contribute to a deeper understanding of the internal regulatory networks underlying the biodegradation mechanism of PentaCB.

## ACKNOWLEDGEMENTS

This work was supported by the National Key Research and Development Program of China (2023YFD1902701); the Shandong Provincial Key Research and Development Program, China (2018GSF117004); the National Natural Science Foundation of China (62102065); the Research Project (Youth Fund) of Shandong Academy of Sciences (2018QN0018); the NSFC-Shandong United Fund (U1906222); the Medical and Health Science and Technology Development Plan of Shandong Province (2017WS473); and the Central-guided Local Science and Technology Development Fund (YDZX2023057).

## SUPPLEMENTARY MATERIALS

**Figure S1.**
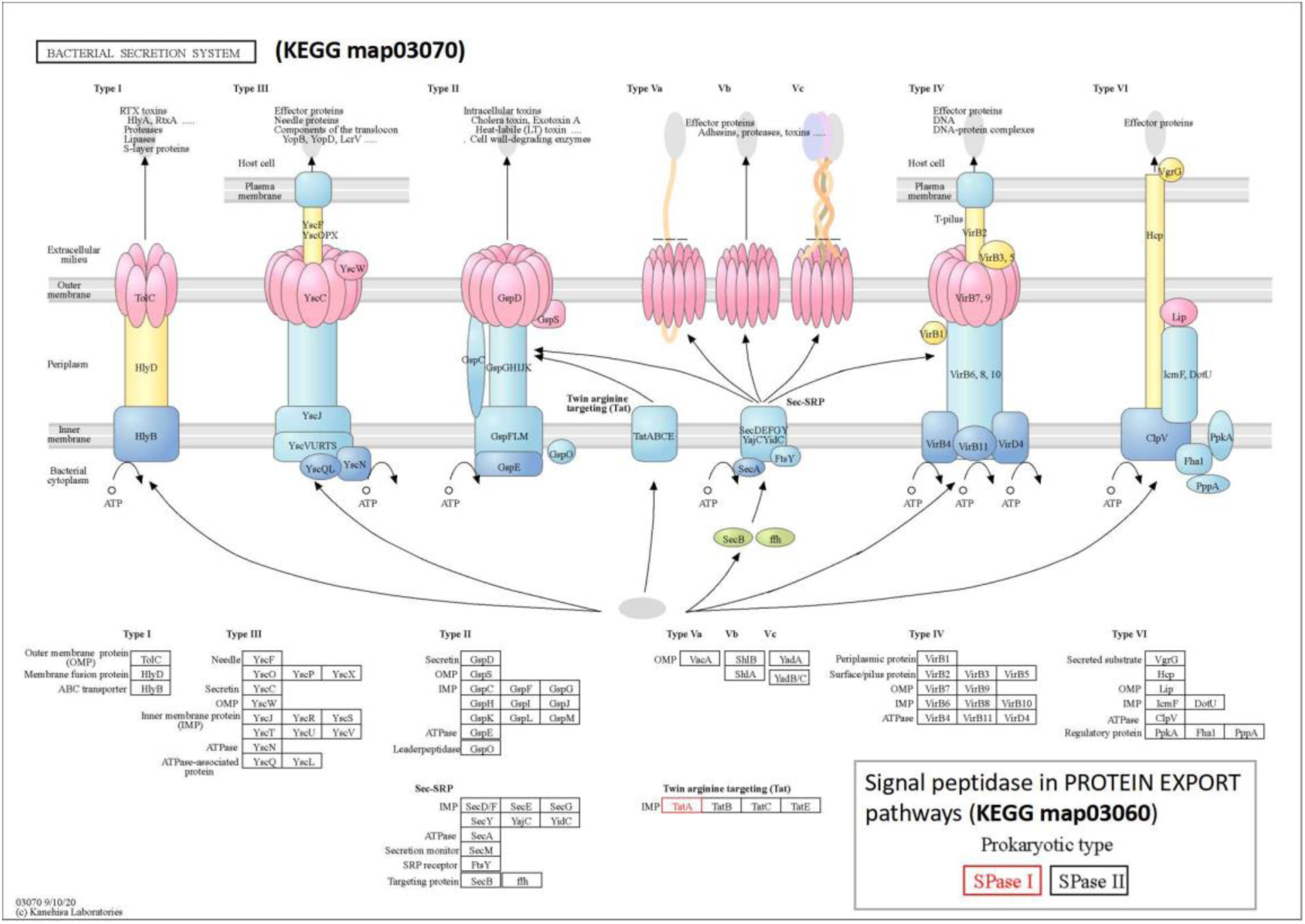
Protein secretory and export pathways were enriched among the up-regulated proteins induced by PentaCB. In the proteomics analysis of *M. paraoxydans* incubated with PentaCB, we detected Signal peptidase I (SPase I), crucial for translocated preprotein release from the cytoplasm to the periplasm, and TatA, a component of the twin-arginine translocation (Tat) system. This suggests that enzymes responsible for PentaCB degradation were distributed both in the cytoplasm and on the outer surface of the cytoplasmic membrane. The image was sourced from the KEGG database, and the up-regulated relevant proteins induced by PentaCB are outlined in red boxes.

**Table 1S.** LB medium.

| Component | Concentration (g·L<sup>-1</sup>) |
| --- | --- |
| Peptone | 10.0 |
| Yeast extract | 5.0 |
| NaCl | 10.0 |

**Table 2S.** Induction medium.

| Component | Concentration (g·L<sup>-1</sup>) |
| --- | --- |
| Peptone | 2 |
| (NH<sub>4</sub>)<sub>2</sub>SO<sub>4</sub> | 1.0 |
| NaCl | 0.2 |
| KH<sub>2</sub>PO<sub>4</sub> | 0.5 |
| K<sub>2</sub>HPO<sub>4</sub> | 0.5 |
| MgSO<sub>4</sub> | 0.2 |
| CaCl<sub>2</sub> | 0.1 |

## S1 TMT Proteomic Quantification and Functional Analysis

### S1.1 Total Protein Extraction

Cells were lysed using liquid nitrogen and then ultrasonication individually. The lysate was centrifugated and the supernatant was collected. The protein concentration was determined using the Bradford protein assay. Extracts were reduced and then alkylated with iodoacetic acid. Samples were precipitated with acetone, collected, and washed with cold acetone. The precipitate was dissolved in buffer, following tests of protein concentration.

### S1.2 Peptide Preparation and TMT Labeling

Supernatants containing 0.1 mg of protein were subjected to Trypsin Gold digestion (enzyme-to-substrate ratio 1:50). Peptides were desalted, dried, and labeled with TMT6/10-plex reagents. After 1 h incubation, the reaction was stopped with ammonium hydroxide. Differentially labeled peptides were combined and desalted. A common reference sample was created from pooling aliquots from each sample.

### S1.3 HPLC Fractionation and LC-MS/MS Analysis

TMT-labeled peptides were fractionated using a C18 column on a Rigol L3000 HPLC with a gradient elution. Eluates were collected and combined into 15 fractions. The fractions were dried under vacuum and reconstituted in 0.1% formic acid (FA). 2 μg peptide samples were introduced into a column, where peptide separation was achieved using a gradient elution method. EASY-nLC 1200 UHPLC system coupled with an Orbitrap Q Exactive HF-X mass spectrometer were used for shotgun proteomics sequencing.

### S1.4 Identification and Quantitation of Protein

Spectra were searched against the *M. paraoxydans* UniProt database using Proteome Discoverer 2.2. Search parameters included mass tolerances, fixed and variable modifications, and miscleavage sites. Protein identification required FDR < 1% on both peptide and protein levels. Proteins with similar peptides, indistinguishable by MS/MS analysis, were grouped separately as protein groups. TMT quantification was performed using Reporter Quantification. The Mann-Whitney Test was used in statistical analysis.

### S1.5 Functional Analysis of Protein and DEP

GO and IPR analyses were performed using the interproscan-5 program against the non-redundant protein database (Pfam, PRINTS, ProDom, SMART, ProSiteProfiles, and PANTHER). Protein family and pathway analysis utilized the COG and KEGG databases. Potential interacting partners were predicted using the STRING-db server (http://string.embl.de/), which contains both known and predicted protein-protein interactions (Franceschini et al., 2013). Enrichment analysis of GO, IPR, and KEGG was performed using an enrichment pipeline (Huang et al., 2008). Differentially expressed proteins (DEP) were identified using significant ratios (p < 0.05 and |log_2_FC| > 1.2).

### S2 UPLC-QTOF-MS Metabolomics Quantification and Analysis S2.1 Metabolites Extraction and GC-MS/MS Analysis

Completely dry the samples using a vacuum concentrator without applying heat. Then, add 60 μL of methoxyamination hydrochloride (20 mg/mL in pyridine) and incubate the mixture at 80°C for 30 min. 80 μL BSTFA reagent (1% TMCS, v/v) was added to the sample aliquots followed by incubation at 70 °C for 1.5 h. GC system coupled with a Pegasus HT TOF-MS was employed for analysis.

### S2.2 Data Analysis

Data processing included peak extraction, baseline correction, peak alignment, deconvolution analysis, peak identification, and area integration were performed using LECO Chroma TOF 4.3X software and the LECO-Fiehn Rtx5 database. Metabolite identification relied on both mass spectrum matching and retention index matching.

